# The Spatial Reach of Neuronal Coherence and Spike-field Coupling across the Human Neocortex

**DOI:** 10.1101/2021.12.07.471617

**Authors:** John C. Myers, Elliot H. Smith, Marcin Leszczynski, James O’Sullivan, Mark Yates, Guy McKhann, Nima Mesgarani, Charles Schroeder, Catherine Schevon, Sameer A. Sheth

**Affiliations:** Department of Neurosurgery, Baylor College of Medicine, Houston, TX; Department of Neurosurgery, University of Utah, Salt Lake City, UT; Department of Psychiatry, Columbia University, New York, NY; Department of Electrical Engineering, Columbia University, New York, NY; Department of Neurology, Columbia University, New York, NY

**Author notes:** Corresponding author John Myers, Ph.D., Baylor College of Medicine, One Baylor Plaza, Houston, TX 77030, Department of Neurosurgery.

**Keywords:** neuronal coherence, spike-field coherence, phase alignment, functional connectivity

## Abstract

Neuronal coherence is thought to be a fundamental mechanism of communication in the brain, where synchronized field potentials coordinate synaptic and spiking events to support plasticity and learning. Although the spread of field potentials has garnered great interest, little is known about the spatial reach of phase synchronization, or neuronal coherence. Functional connectivity between different brain regions is known to occur across long distances, but the locality of coherence within a brain region is understudied. Here we used simultaneous recordings from electrocorticography (ECoG) grids and high-density microelectrode arrays to estimate the spatial reach of neuronal coherence and spike-field coherence (SFC) across frontal, temporal, and occipital cortices during cognitive tasks in humans. We observed the strongest coherence within a 2-3 cm distance from the microelectrode arrays, potentially defining an effective range for local communication. This range was relatively consistent across brain regions, spectral frequencies, and cognitive tasks. The magnitude of coherence showed power law decay with increasing distance from the microelectrode arrays, where the highest coherence occurred between ECoG contacts, followed by coherence between ECoG and deep cortical LFP, and then SFC (i.e., ECoG > LFP > SFC). The spectral frequency of coherence also affected its magnitude. Alpha coherence (8-14 Hz) was generally higher than other frequencies for signals nearest the microelectrode arrays, whereas delta coherence (1-3 Hz) was higher for signals that were farther away. Action potentials in all brain regions were most coherent with the phase of alpha oscillations, which suggests that alpha waves could play a larger, more spatially local role in spike timing than other frequencies. These findings provide a deeper understanding of the spatial and spectral dynamics of neuronal coherence, further advancing knowledge about how activity propagates across the human brain.

## INTRODUCTION

The human brain can remarkably extract rich sensory information from the outside world and quickly integrate it with higher order cognitive processes. Abilities such as controlled decision making, speech perception, and social interaction all require interactions between spatially distributed action potentials and neural oscillations across the brain (Meunier et al., 2009; Perlovsky, 2013; Smith et al., 2019). Field potentials (FP) are the electric fields of the brain, reflecting transmembrane currents within neural tissue (Buzsáki et al., 2012; Eccles, 1951; Galindo-Leon and Liu, 2010; Kajikawa and Schroeder, 2015; Mitzdorf, 1985; Womelsdorf et al., 2007). The spatial spread of FPs has become of great interest in neuroscience due to its fundamental influence on neural computations and behavior (Kajikawa and Schroeder, 2011; Katzner et al., 2009; Lindén et al., 2011; Xing et al., 2009). FPs can be measured with surface electrodes or intraparenchymal probes, but the material, spacing, and impedance of the electrodes can potentially affect the estimated spatial extent (Buzsáki et al., 2012; Dubey and Ray, 2019; Lee et al., 2005). Previous studies have shown that FPs can be confined to very small domains (200-400 *μ*m) under tightly controlled conditions, such as when stimuli are confined to single unit receptive fields in anesthetized monkeys (Xing et al., 2009). When stimuli are embedded in complex naturalistic scenes that excite ensembles of neurons in awake monkeys, FPs can reach many millimeters and even centimeters from a current source (Dubey and Ray, 2019; Kajikawa and Schroeder, 2011). During epileptic seizures, multiunit spiking activity can organize into low frequency oscillations detectable within 10 cm from the onset zone (Eissa et al., 2017). Thus there is evidence to suggest that FPs may reach much farther than would normally be considered ‘local.’ Empirical and computational models suggest that spatial reach can also vary with frequency, synaptic distribution, and neuronal morphology (Dubey and Ray, 2016; Leski et al., 2013; Lindén et al., 2011; Rasch et al., 2008; Kajikawa and Schroeder 2015).

The phase of FPs has been linked to cognitive processes such as perception, attention, and decision making (Bosman et al., 2012; Busch et al., 2009; Lakatos et al., 2005; 2008; Leszczynski and Schroeder, 2019; Mathewson et al., 2009; Neuling et al., 2012; Solomon et al., 2017; Tal et al., 2020; Womelsdorf and Fries, 2006). Coherent FPs can facilitate the transfer of information across shorter and longer distances (Fries, 2005; Helfrich and Knight, 2016; Lachaux et al., 1999; Liebe et al., 2012; Rodriguez et al., 1999; Varela et al., 2001). Neuronal coherence has been shown to support higher order cognitive processes (Benchenane et al., 2010; Fries, 2015; Lachaux et al., 1999; Oehrn et al., 2014; Rodriguez et al., 1999; Varela et al., 2001). Therefore, measuring the spatial reach of coherence is an important step towards understanding the nature of rhythmic information flow across the brain.

We had a rare opportunity to measure both superficial and deep cortical FPs from simultaneous ECoG and microelectrode array (MEA) recordings in three human neurosurgical patients undergoing intracranial epilepsy monitoring. The MEAs allowed us to measure action potentials from nearly 500 neurons, most likely located in layers 4/5 (Schevon et al., 2012). Our primary goal was to estimate the spatial reach of surface and depth FP coherence and spike-field coherence (SFC) in the human neocortex. Another key goal was to determine how the properties of coherence might change based on oscillatory frequency and distance from the MEA. We hypothesized that coherence would decrease with distance from the point of origin, and that the frequencies of highest coherence would vary with tissue depth (superficial cortical FP vs. deep cortical FP).

## RESULTS

We measured neuronal coherence between field potentials using data from three neurosurgical patients who underwent seizure monitoring for drug-resistant epilepsy. Each patient had subdural ECoG grids placed in a different brain region, along with a Utah microelectrode array to simultaneously record deep cortical FP and action potentials from a total of 497 neurons. The brain regions covered were the frontocentral cortex (8 × 8 contacts, 0.5 cm spacing; 100 single units), middle temporal cortex (8 × 8 contacts, 1 cm spacing; 191 single units), and posterior temporal/occipital cortex (8 × 3 contacts, 1 cm spacing; 206 single units; Figure 1). The subject with frontocentral grid coverage (subject A) performed a controlled decision-making task, the subject with middle/superior temporal coverage (subject B) performed a speech perception task, and the subject with posterior temporal/occipital coverage (subject C) performed a social cognition task. Each task was intended to evoke some degree of region-specific activity (see **METHODS** for task descriptions).

**Figure 1.**
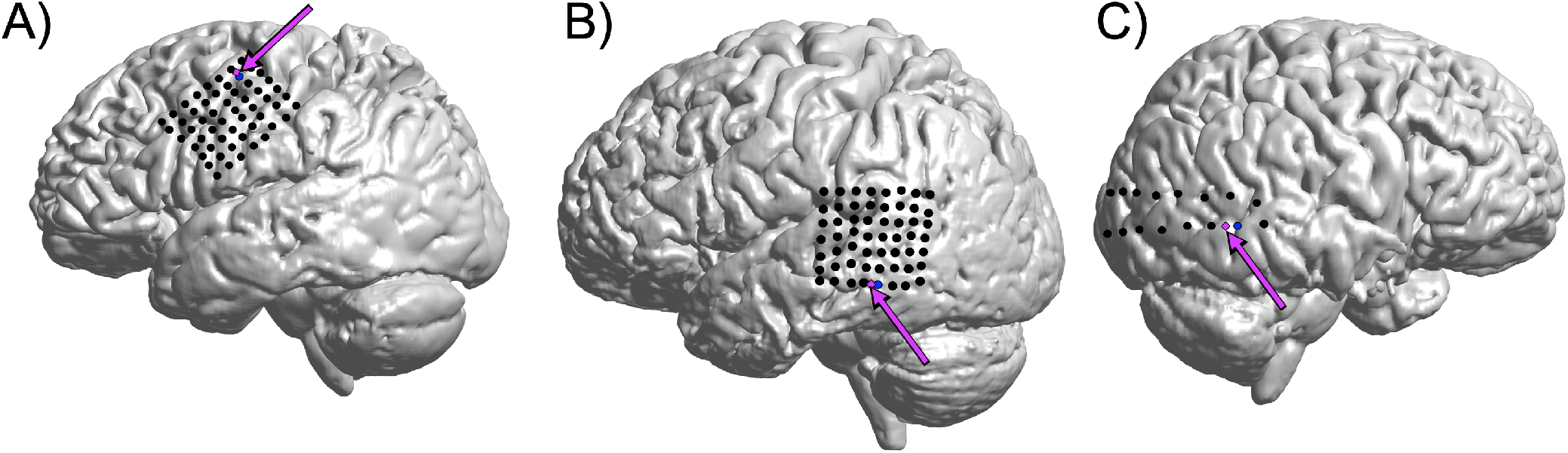
ECoG Grids and Microelectrode Arrays. Each subject (A-C) had a subdural ECoG grid placed in the frontocentral (subject A), middle temporal (subject B), or posterior temporal cortex (subject C), respectively. Each subject was also implanted with a microelectrode array (MEA) (magenta arrows) that allowed recording of deep cortical FP and action potentials from single neurons. The location of the ECoG channel closet to the MEA (blue) was treated as the spatial origin for computing distance across the grid.

### Coherence is Inversely Proportional to Distance across the Neocortex

The estimated spatial reach of coherence across the cortical surface was computed by first modeling coherence as a function of Euclidean distance (mm) from the microelectrode array (MEA) in each brain region. Change-point detection (CPD) was used to estimate spatial reach by finding the distance at which variance in the magnitude of coherence changed significantly (see **METHODS** for CPD description). The propagation of neural signals across the cortical mantle is thought to conform to power law dynamics, where the sum or magnitude of neural signals are inversely proportional to distance from their origin (Klaus et al., 2011). We found that coherence decreased as a power function of distance from the microelectrode arrays (Figure 2). To investigate interaction effects between distance and oscillatory frequency, we computed a median split across all contacts based on the median distance from the MEA (2.29 cm). The term ‘proximal coherence’ refers to coherence with signals within the median distance from the MEA (< 2.29 cm). ‘Distal coherence’ refers to coherence with contacts farther away from the MEA (> 2.29 cm).

**Figure 2.**
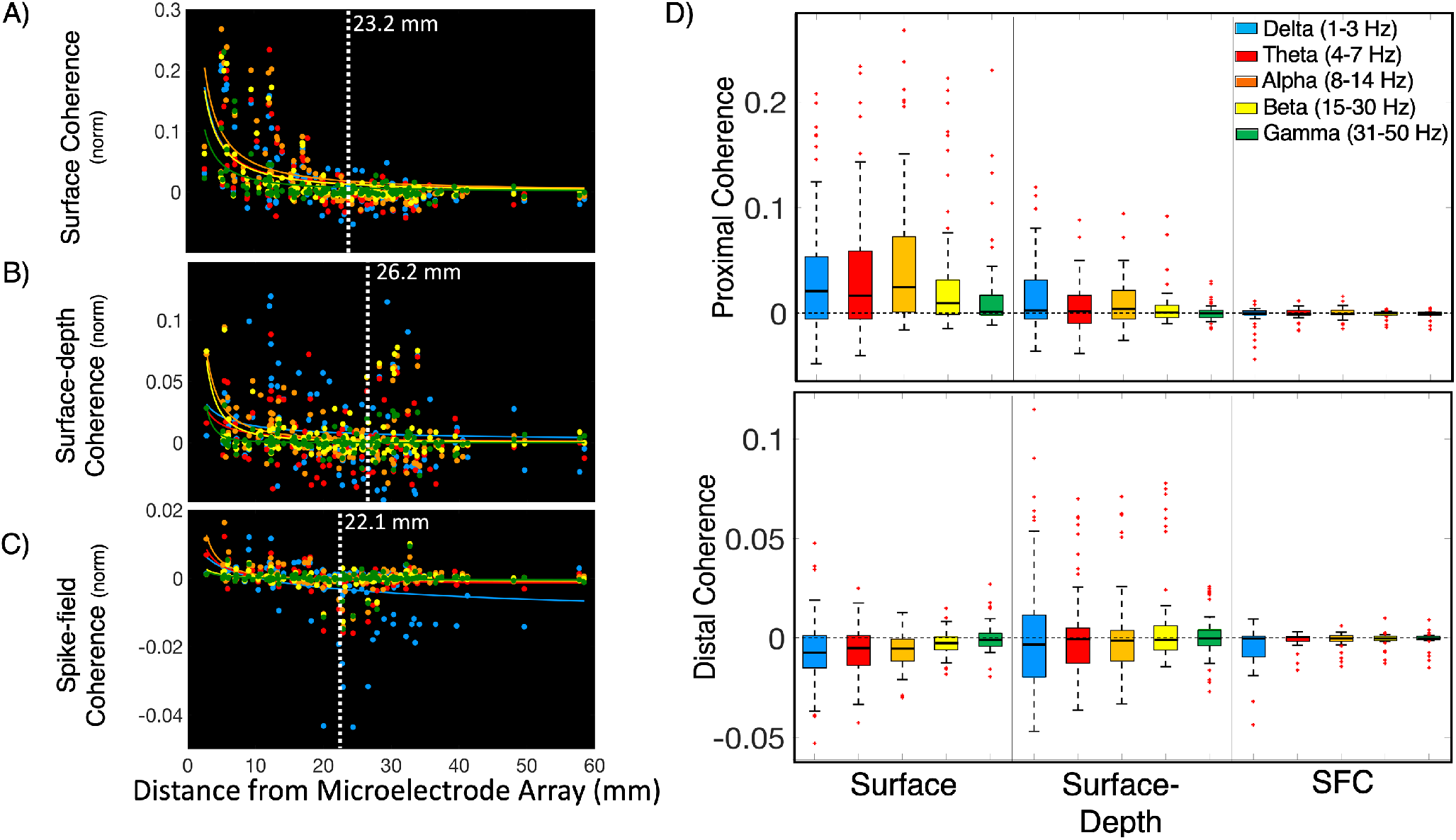
Coherence across Brain Regions Decreases as a Function of Distance. (A) Across frontal and temporal cortices, neuronal coherence between ECoG contacts decreased with distance. Power law decay of surface coherence was observed in delta, theta, alpha, beta, and gamma frequency bands (*p* < 0.001). (B) Neuronal coherence between surface (ECoG) and deep cortical field potentials (MEA) also decreased with distance for alpha (*p* < 0.001), beta (*p* < 0.001), and gamma frequencies (*p* < 0.001). (C) Spike-field coherence (SFC) between action potentials and superficial FPs (ECoG) showed a similar pattern of power law decay for delta (*p* = 0.003), theta (*p* < 0.001), and alpha (*p* < 0.001). Vertical white lines depict the spatial reach of coherence, which was estimated using changepoint detection on the coherence x distance data across all brain regions. (D) ‘Proximal Coherence’ refers to ECoG contacts that were within the median distance of the microelectrode array (2.29 cm), whereas ‘Distal Coherence’ refers to contacts that were beyond that range. The magnitude of surface coherence was greater for proximal contacts, especially for alpha signals (*p* < 0.001). Delta and theta coherence between superficial and deep layer field potentials did not decrease significantly with distance. The results reported in A, B, and C were computed by fitting an power function to the median-centered coherence data as a function of distance from the microelectrode arrays. White vertical lines indicate the estimated spatial reach based on CPD. P-values indicate the significance of correlation between the raw and predicted values of coherence as a function of distance. Values above the black horizontal lines in D represent channels where coherence was greater than the median value. The results shown in D were tested with a 3 (depth: surface, surface-depth, SFC) x 5 (*frequency*: delta, theta, alpha, beta, gamma) x 2 (*distance*: proximal, distal) ANOVA.

We examined 125 ECoG signals from frontocentral (n = 55), middle temporal (n = 55), and temporo-occipital cortices (n = 15). Pooled all brain regions, coherence between ECoG contacts, or ‘surface coherence’, decreased with distance from the microelectrode arrays. CPD estimated that the spatial reach of surface coherence was 2.32 cm. Significant power law decay of surface coherence was observed for delta (1-3 Hz) (*r*^*2*^ = 0.286, *p* < 0.001), theta (4-7 Hz) (*r*^*2*^ = 0.25, *p* < 0.001), alpha (8-14 Hz) (*r*^*2*^ = 0.32, *p* < 0.001), beta (15-30 Hz) (*r*^*2*^ = 0.30, *p* < 0.001), and gamma bands (31-50 Hz) (*r*^*2*^ = 0.24, *p* < 0.001) (Figure 2A). The *r*^*2*^ values indicate the goodness-of-fit and significance of the correlations between the raw and model-predicted values of coherence as a function of distance. We then used a 2 (*distance*: proximal, distal) x 5 (*frequency*: delta, theta, alpha, beta, gamma) ANOVA to test how distance and frequency affected the magnitude of surface coherence. A main effect of distance showed that surface coherence was greater for contacts within a 2.29 cm median distance around the MEA, *F*(1, 62) = 35.08, p *<* 0.001, *η*^*2*^ = 0.36. Surface coherence in the theta (4-7 Hz) and alpha bands (8-14 Hz) was higher than other frequencies, *F*(4, 59) = 4.22, p *=* 0.005, *η*^*2*^ = 0.22. However, a distance x frequency interaction showed that theta and alpha bands only had the highest coherence for signals within the median distance, *F*(4, 59) = 8.86, p *<* 0.001, *η*^*2*^ = 0.38. Surface coherence in the gamma band was higher than other frequencies for ECoG channels outside the median distance, *F*(4, 59) = 3.376, p *<* 0.001, *η*^*2*^ = 0.336 (Figure 2D). These findings suggest that the phase alignment of superficial FPs spreads across several millimeters of the same brain region with a certain degree of frequency specialization. Higher gamma coherence may indicate temporally aligned inputs to superficial cortical layers (Buzsáki and Schomburg, 2015).

Coherence between superficial (ECoG) and deep cortical FPs recorded from the MEA will be referred to as ‘surface-depth coherence’. CPD estimated that the spatial reach of surface-depth coherence was 2.62 cm from the MEAs, which is approximately 3 mm farther than surface coherence. Surface-depth coherence showed power law decay with distance for alpha (*r*^*2*^ = 0.17, *p* < 0.001), beta (*r*^*2*^ = 0.08, *p* < 0.001), and gamma frequency bands (*r*^*2*^ = 0.09, *p* < 0.001). Delta (*p* = 0.054) and theta (*p* = 0.0502) surface-depth coherence did not show power law decay across distance (Figure 2B). Across all frequencies, the magnitude of surface-depth coherence did not decrease with distance from the MEA (p = 0.142). Delta surface-depth coherence (1-3 Hz) seemed to overshadow other frequencies, but only for signals within the median distance, *F*(4, 59) = 5.16, p *=* 0.001, *η*^*2*^ = 0.26. Signals outside the median distance were more coherent within the beta band (15-30 Hz), *F*(4, 59) = 2.76, p *=* 0.036, *η*^*2*^ = 0.16. These findings are convergent with previous research emphasizing the importance of delta and beta oscillations for cross-laminar communication (Bastos et al., 2018; Sotero et al., 2015) (Figure 2B).

Spike-field coherence (SFC) between action potential spike trains and ECoG reached approximately 2.21 cm from the MEAs, which was the shortest distance across all types of coherence that we measured (Figure 2C). We observed significant power law decay for delta (*r*^*2*^ = 0.07, *p* = 0.003), theta (*r*^*2*^ = 0.12, *p* = 0.001), and alpha (*r*^*2*^ = 0.20, *p* < 0.001) SFC. Beta (*p* = 0.056) and gamma SFC (*p* = 0.164) did not show power law decay. SFC did not decrease significantly with outside the median radius but it trended downwards (p = 0.073). A distance x frequency interaction revealed that alpha SFC was higher for proximal signals, whereas beta and gamma SFC were higher for distal signals, *F*(4, 58) = 2.97, p *=* 0.027, *η*^*2*^ = 0.17 (Figure 2D).

Across all types of coherence (i.e., surface, surface-depth, SFC), a main effect of depth showed that surface coherence was higher than surface-depth and SFC, *F*(2, 61) = 19.41, p *<* 0.001, *η*^*2*^ = 0.389 (Figure 2D). This result implies more synchrony among neurons in within layers than across layers. Intriguingly, surface-depth coherence was higher than surface coherence for signals beyond the median distance, which is convergent with the slightly higher estimates of spatial reach (23 mm vs. 26 cm), *F*(2, 61) = 17.46, p *<* 0.001, *η*^*2*^ = 0.36 (Figure 2B). These results suggest that coherence involving deeper cortical layers has the potential to reach across similar distances to coherence across superficial layers. Overall, alpha coherence was higher than other frequencies across all types of coherence (*η*^*2*^ = 0.243, p *<* 0.001). The prominence of alpha synchrony may relate to a specialized role in cortico-cortical communication (Chapeton et al., 2019).

### Frontocentral Cortical Coherence

Subject A had ECoG grid coverage (n = 55 signals) across the frontocentral cortex, including the middle frontal, precentral, and postcentral gyri (Figure 1A, 3). These brain regions play important roles in cognitive control and response selection. Subject A performed the multi-source interference task (MSIT), which is a controlled decision-making task with two dimensions of decision conflict (Sheth et al., 2012; Smith et al., 2019) (see **METHODS** for task descriptions). The microelectrode array was placed on the posterior middle frontal gyrus (MFG). Across frontocentral cortex, surface coherence reached an estimated 2.85 cm, showing significant power law decay for delta (*r*^*2*^ = 0.38, *p* < 0.001), theta (*r*^*2*^ = 0.40, *p* < 0.001), alpha (*r*^*2*^ = 0.45, *p* < 0.001), beta (*r*^*2*^ = 0.45, *p* < 0.001), and gamma (*r*^*2*^ = 0.39, *p* < 0.001) signals (Figure S1A). A main effect of distance showed that surface coherence was greater for contacts within the median distance of 2.22 cm around the MEA, *F*(1, 27) = 20.85, p *<* 0.001, *η*^*2*^ = 0.44. Delta (1-3 Hz) and theta band (4-7 Hz) coherence were most prominent across the surface of frontal cortex, *F*(4, 24) = 234.93, p *<* 0.001, *η*^*2*^ = 0.98. Delta coherence was higher than other frequencies for signals outside the median distance, which may implicate the role of slower oscillations in distal communication, *F*(4, 24) = 7.83, p *<* 0.001, *η*^*2*^ = 0.566 (Figure 6A).

**Figure 3.**
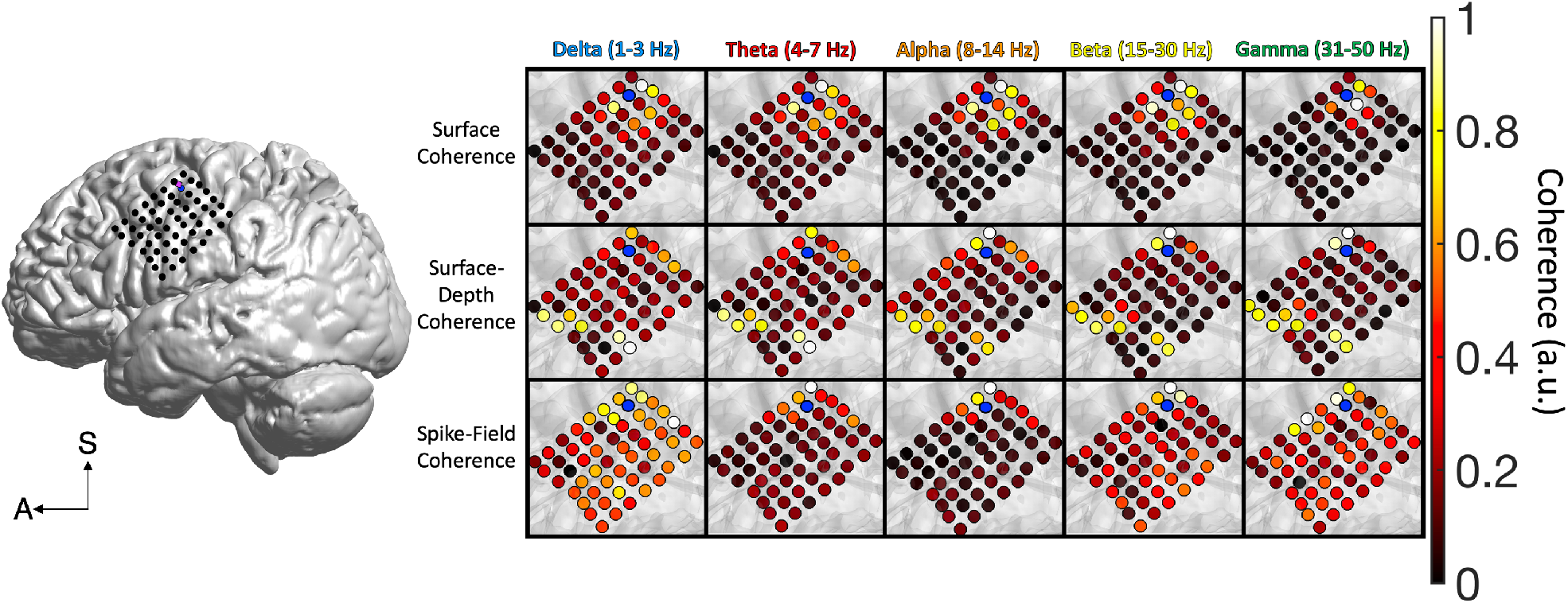
Frontocentral Cortical Coherence. The spatial spread of surface coherence (ECoG-to-ECoG), surface-depth coherence (ECoG-to-FP), and spike-field coherence (SFC) across frontal cortex. The ECoG channel closest to the microelectrode array (blue) was treated as the spatial origin point for distance computations. The data in this figure were max-min normalized (0-1) to highlight the spatial patterns of each frequency band and type of coherence. legend in the lower left corner defines the superior (S) and anterior (A) axes.

**Figure 4.**
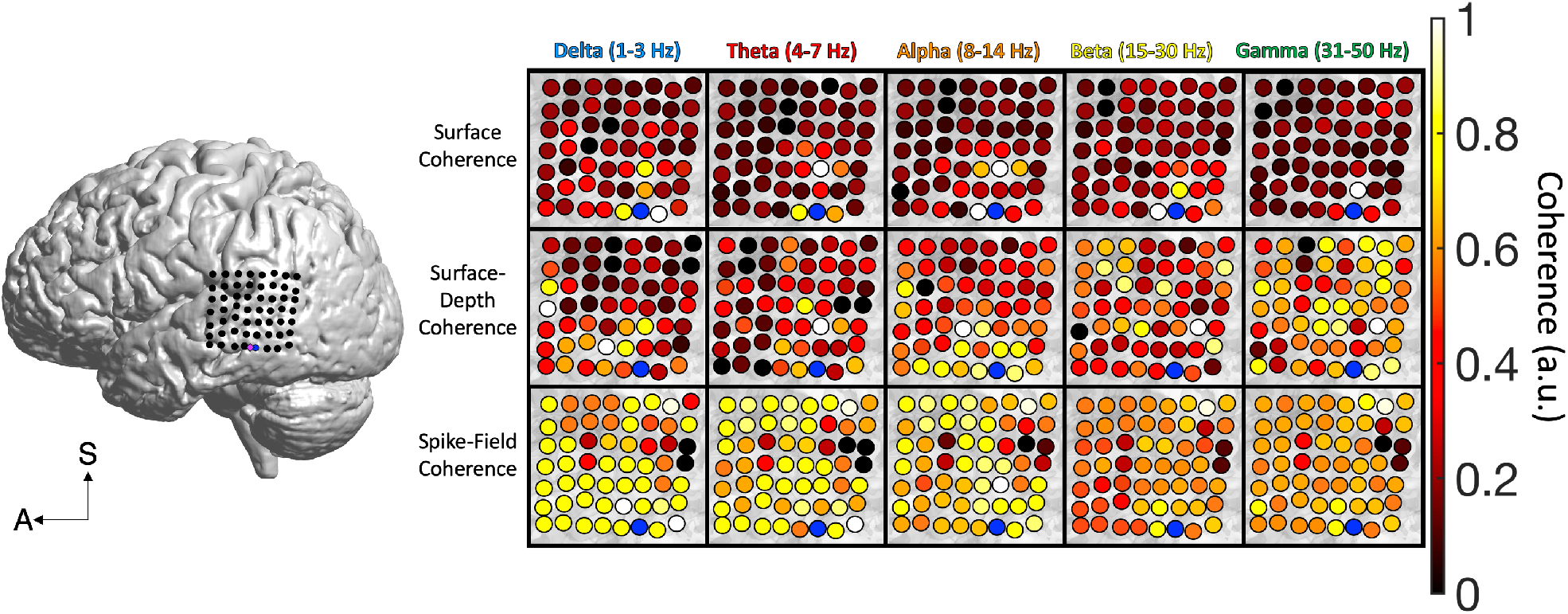
Middle Temporal Coherence. The spatial spread of surface coherence, surface-depth coherence, and SFC across middle temporal cortex. The ECoG channel closest to the microelectrode array (blue) was treated as the spatial origin point for distance computations. The data in this figure were max-min normalized (0-1) to display the

**Figure 5.**
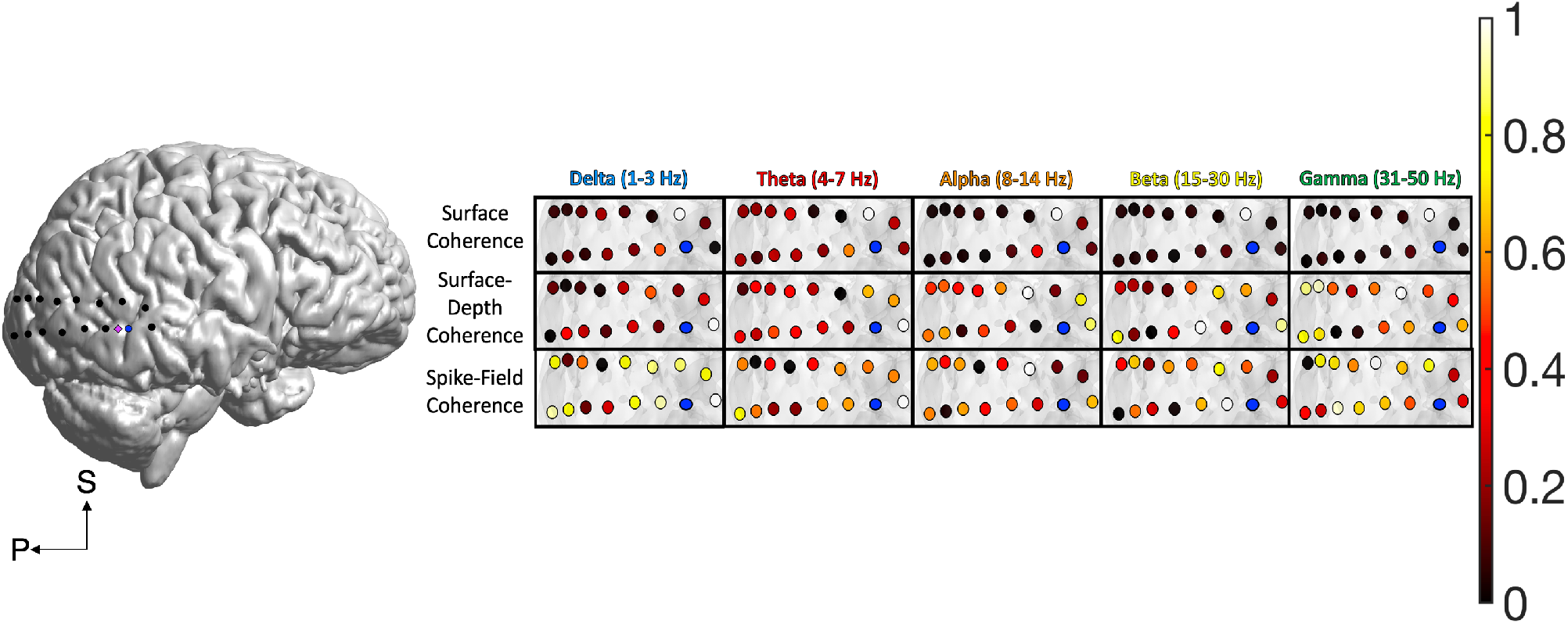
Posterior Temporal and Occipital Coherence. The spatial spread of surface coherence, surface-depth coherence, and spike-field coherence (SFC) across posterior temporal cortex. The ECoG channel closest to the microelectrode array (blue) was treated as the spatial origin point for distance computations. The data in this figure were max-min normalized (0-1) to highlight the spatial patterns of each frequency band and type of coherence. The legend in the lower left corner defines the superior (S) and anterior (A) axes.

**Figure 6.**
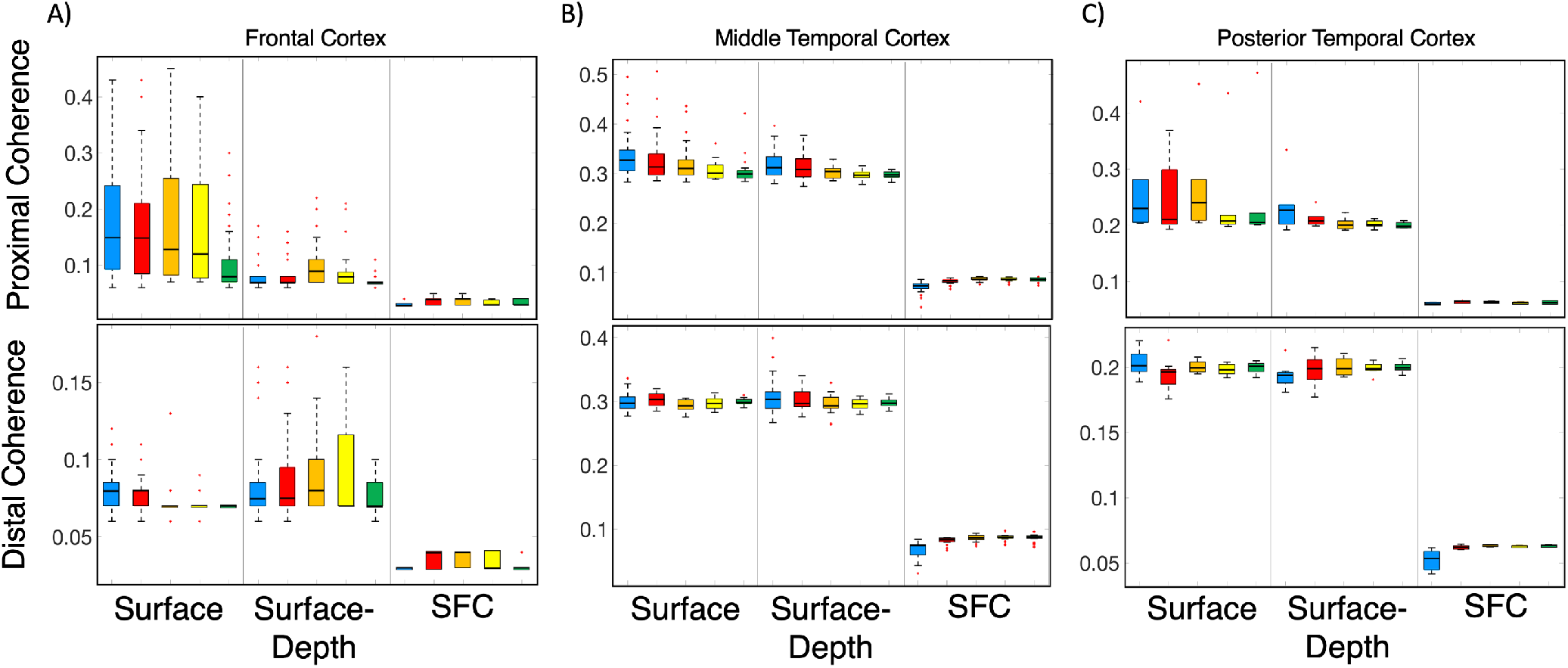
Proximal vs. Distal Coherence for Frontal and Temporal Cortices. Across the frontocentral cortex, alpha signals had the highest magnitude of surface coherence, particularly for proximal channels (p *<* 0.001). Delta surface coherence was higher for distal channels (p *=* 0.037). Alpha signals in frontocentral cortex showed the highest surface-depth coherence (p *<* 0.001) and SFC as well (p *<* 0.001). Across the middle temporal cortex, the magnitude of surface coherence was highest for delta and theta coherence (p *=* 0.010). Delta surface-depth coherence had the highest magnitude (p *=* 0.001), but delta SFC was the lowest magnitude. Across the posterior temporal cortex, alpha surface coherence was highest for proximal channels, and delta coherence was highest for distal channels (p *=* 0.033). Theta surface-depth coherence was highest for distal channels (p < 0.001), and the magnitude of delta SFC fell off significantly with distance across posterior temporal/occipital regions (p *=* 0.024). These results were computed using 2 (*distance*: proximal, distal) x 3 (*depth*: surface, surface-depth, SFC) 5 (*frequency*: delta, theta, alpha, beta, gamma) ANOVAs.

The estimated spatial reach of surface-depth coherence across the frontocentral cortex was 3.67 cm (Figure S1A). Across all frequencies, the magnitude of surface-depth coherence was comparable for proximal and distal signals, showing no substantial decreases beyond the median distance (*p* = 0.698). Surface-depth coherence showed power law decay only for alpha oscillations (*r*^*2*^ = 0.09, *p* = 0.027). Delta (*p* = 0.829), theta (*p* = 0.876), beta (*p* = 0.182) and gamma (*p* = 0.394) coherence did not show patterns of decay. Delta and alpha surface-depth coherence were higher within the median distance, but delta and theta coherence were more prominent beyond the median distance, *F*(4, 24) = 4.00, p *=* 0.013, *η*^*2*^ = 0.40. These results suggest that surface-depth coherence has the potential to retain its magnitude across a greater distance than surface coherence (Figure 6A).

Spike-field coherence (SFC) was averaged across 100 simultaneously recorded neurons in the posterior middle frontal gyrus (MFG). SFC reached an estimated 2.77 cm across the frontocentral cortex. Power law decay with distance was observed for delta (*r*^*2*^ = 0.12, *p* = 0.011), theta (*r*^*2*^ = 0.39, *p* < 0.001), alpha (*r*^*2*^ = 0.41, *p* < 0.001), beta (*r*^*2*^ = 0.16, *p* = 0.002), and gamma SFC (*r*^*2*^ = 0.246, *p* = 0.001). SFC decreased significantly beyond the median distance, *F*(1, 27) = 4.43, p *=* 0.045, *η*^*2*^ = 0.14. The magnitude of SFC was greatest for alpha signals, across both proximal and distal contacts, *F*(4, 24) = 267.48, p *<* 0.001, *η*^*2*^ = 0.98 (Figure 6A). Action potentials recorded from the MFG were more coherent with the phase of alpha oscillations.

Across all types of coherence in frontocentral cortex, a main effect of depth showed that surface coherence was higher than surface-depth coherence and SFC, *F*(2, 26) = 4040.99, p *<* 0.001, *η*^*2*^ = 0.99 (Figure 2). However, a distance x depth interaction revealed that beyond the median distance, surface-depth coherence was greater than surface coherence, *F*(2, 26) = 10.10, p *=* 0.001, *η*^*2*^ = 0.44 (Figure 6A). A main effect of frequency showed that delta and theta coherence were highest across all types of coherence in frontocentral cortex [*F*(4, 24) = 292.86, p *<* 0.001, *η*^*2*^ = 0.98], but a depth x frequency interaction showed that the dominant frequency bands were unique to each depth, *F*(8, 20) = 451.84, p *<* 0.001, *η*^*2*^ = 0.99. Delta and theta coherence were more prominent for surface coherence. Surface-depth coherence was dominated by delta and alpha frequencies, whereas theta and alpha dominated SFC. These results suggest that phase synchrony within frontocentral cortex has nuanced patterns that could arise from tissue depth and oscillatory frequency.

### Middle and Superior Temporal Coherence

Subject B had ECoG coverage across the temporal cortex, including the middle, superior, and inferior temporal gyri, and the supramarginal gyrus. These regions are known to play important roles in auditory perception and attention. The participant completed a speech perception task in which they listened to two speakers (male and female) talking at the same time. The goal was to attend to either the male or female speaker in alternating blocked trials (O’Sullivan et al., 2019) (see **METHODS** for task descriptions). The microelectrode array was placed on the middle temporal gyrus (MTG). Surface coherence reached approximately 2.44 cm, showing power law decay for delta (*r*^*2*^ = 0.48, *p* < 0.001), theta (*r*^*2*^ = 0.33, *p* < 0.001), alpha (*r*^*2*^ = 0.38, *p* < 0.001), beta (*r*^*2*^ = 0.48, *p* < 0.001), and gamma (*r*^*2*^ = 0.21, *p* < 0.001) signals (Figure S1B). A main effect of distance showed significant reductions in surface coherence beyond the median distance of 2.29 cm, *F*(4, 24) = 14.79, p *=* 0.001, *η*^*2*^ = 0.35. Delta coherence was most prominent across the surface of temporal cortex, *F*(4, 23) = 4.305, p *=* 0.010, *η*^*2*^ = 0.428 (Figure 6B).

Surface-depth coherence reached 2.43 cm across temporal cortex, decreasing significantly beyond the median distance *F*(4, 24) = 6.292, p *=* 0.001, *η*^*2*^ = 0.523. We observed power law decay for delta (*r*^*2*^ = 0.13, *p* = 0.009), theta (*r*^*2*^ = 0.09, *p* = 0.022), and alpha signals (*r*^*2*^ = 0.37, *p* = 0.001). Beta (*r*^*2*^ = 0.008, *p* = 0.506), and gamma (*r*^*2*^ = 0.041, *p* = 0.137) surface-depth coherence did not have significant power law decay (Figure S1B). Delta and theta surface-depth coherence were highest across the MTG/STG, *F*(4, 24) = 13.96, p *<* 0.001, *η*^*2*^ = 0.69.

Spike-field coherence (SFC) was averaged across 191 simultaneously recorded neurons recorded in the MTG, reaching an estimated 2.27 cm across temporal cortex. SFC did not significantly reduce beyond the median distance (*p* = 0.368), and significant power law decay was only observed for delta (*r*^*2*^ = 0.095, *p* = 0.022) SFC. Theta (*p* = 0.219), alpha (*p* = 0.128), beta (*p* = 0.239) and gamma SFC (*p* = 0.397) showed non-significant power law decay (Figure S1B). The higher frequencies (i.e., alpha, beta, and gamma) showed greater SFC across the MTG/STG, which suggests tighter coupling between action potentials and higher frequency signals in this region, *F*(4, 24) = 66.47, p *<* 0.001, *η*^*2*^ = 0.92.

Across all types of coherence in MTG/STG cortex, surface coherence was higher than surface-depth coherence and SFC, *F*(2, 26) = 6723.41, p *<* 0.001, *η*^*2*^ = 0.99 (Figure 6B). However, for signals outside the median distance, surface-depth coherence was higher than other forms of coherence, *F*(2, 26) = 9.10, p *=* 0.001, *η*^*2*^ = 0.42 (Figure 6B). This finding aligned with the results from Subject A. Coherence involving deeper cortical layers may have a broader spatial range than superficial layers. Lower frequency (delta-theta) coherence was most prominent for both surface and surface-depth types, and SFC was greater for higher frequencies (alpha, beta, gamma), *F*(8, 20) = 26.99, p *<* 0.001, *η*^*2*^ = 0.915. These findings suggest that coherence across the temporal cortex had a slower rhythm (1-7 Hz), while the time course of action potentials were more aligned with bands of higher frequency components (8-50 Hz).

### Posterior Temporal and Occipital Coherence

Subject C had ECoG coverage across temporal and occipital cortices, including the middle and superior temporal gyri, and the inferior, middle, and superior occipital gyri (see Figure 1C). These brain areas are important for memory and visual perception. Subject C completed a social cognition task in which they watched short video clips from popular movies before answering questions about the semantic and emotional content of the videos (see **METHODS** for task descriptions). The microelectrode array was placed in posterior temporal cortex. Surface coherence reached approximately 2.32 cm and did not decrease significantly beyond the median distance of 3.03 cm (*p* = 0.254). Across all types of coherence in posterior temporal/occipital cortex, surface coherence was higher magnitude than surface-depth coherence and SFC across the posterior temporal cortex, *F*(2, 14) = 256.54, p < 0.001, *η*^*2*^ = 0.97 (Figure 6C). Although surface coherence retained similar magnitudes across distance, we observed showed power law decay for delta (*r*^*2*^ = 0.35, *p* = 0.020), theta (*r*^*2*^ = 0.28, *p* = 0.043), alpha (*r*^*2*^ = 0.50, *p* = 0.003), beta (*r*^*2*^ = 0.31, *p* = 0.031), and gamma (*r*^*2*^ = 0.33, *p* = 0.025) coherence. Delta and alpha were the primary rhythms of coherence across the surface of posterior temporo-occipital cortex, *F*(4, 28) = 5.80, p *=* 0.002, *η*^*2*^ = 0.453.

Surface-depth coherence reached an estimated 2.61 cm, and showed no power law decay with distance from the MEA (*p* > 0.08). Delta (1-3 Hz) was the most prominent rhythm of surface-depth coherence, *F*(4, 28) = 7.36, p *<* 0.001, *η*^*2*^ = 0.51. (Figure 6C, Figure S1C). Spike-field coherence (SFC) was averaged across 206 simultaneously recorded neurons recorded in posterior temporal cortex, reaching an estimated 2.21 cm. SFC did not show significant power law decay in this region (*p* > 0.05). A marginally significant effect of distance showed lower SFC for signals beyond the median distance of 3.03 cm (p = 0.067, *η*^*2*^ < 0.40). A main effect of frequency indicated that spike trains were most coherent with alpha oscillations, which was consistent with the effects observed in subjects A and B, *F*(4, 28) = 40.37, p *<* 0.001, *η*^*2*^= 0.85.

## DISCUSSION

The fundamental relationship between field potentials, single unit activity, and behavior has evoked much curiosity about the spatial reach of the FP signal (Kajikawa & Schroeder, 2011; 2015; Katzner et al., 2009; Leszczynski et al., 2020; Lindén et al., 2011; Xing et al., 2009). Previous studies on the spatial reach of FPs focused on brain regions with unique cytoarchitecture such as visual (Katzner et al., 2009) or auditory cortex in non-human primate models (Kajikawa and Schroeder, 2011, 2015; Smith et al., 2013). There remains a significant gap in our knowledge about the spatial properties of FPs in the human brain. We used simultaneous ECoG and high-density UMA recordings to estimate the spatial reach of neuronal coherence and SFC across the human neocortical surface. Our methods provide insight about the distance that neuronal coherence might reach across a given brain region. Across frontocentral and temporal cortices in humans, coherence reached 2-3 cm. Our findings were consistent with evidence showing that FPs are detectable across several millimeters (Kajikawa and Schroeder, 2011). By studying coherence specifically, we have quantified the spatial reach of phase alignment itself, which seems to be directly involved in neuronal communication because phase synchronization can facilitate coordinated neuronal activity (Fries, 2015; Lachaux et al., 1999). The spatial reach of coherence was similar across brain regions and tasks, which suggests that these findings may generalize to neurophysical principles. Understanding these spatial properties may shed light on the nature of local neuronal communication and the spread of synchronous activity across the neocortex.

Previous reports have shown that the tips of UMA electrodes (1.5 mm) can record from layers 4/5 of the cortex (Schevon et al., 2012), which reinforced our confidence in exploring the lateral spread of ‘surface-depth’ coherence. The oscillatory and single unit responses to different stimulus inputs are known to vary within brain regions. For example, neurons in layer 4 of auditory cortex will respond in classic feedforward fashion to auditory stimuli, but tactile stimuli elicit responses in more the superficial, supragranular layer (Lakatos et al., 2007). A related physiological mechanism could be thalamocortical calbindin-positive matrix neurons, which begin in the thalamus and project to superficial cortical layers (Jones, 2001). These matrix neurons could play a key role in corticocortical coherence (Müller et al., 2020; Schroeder and Lakatos, 2009). Our results may link to these previous observations, implying that longer range cortico-cortical synchrony can spread via deeper cortical layers. Surface-depth coherence was greater for distances beyond 2.3 cm, peaking for delta and theta signals.

Across all brain regions, the magnitude of SFC was greatest for alpha signals (8-14 Hz), which could highlight a nascent link between alpha and local spike timing in neocortex. Some theorize that alpha oscillations play a specific role in excitatory-inhibitory balance (Chapeton et al., 2019; Klimesch et al., 2007; Mathewson et al., 2011). Oscillatory timing of neuronal activity allows for complex coding schemes such as ‘temporal coding,’ where neural populations synchronize to process complex stimuli (Engel et al., 1991; Panzeri et al., 2001; Kayser et al., 2009; Panzeri et al., 2010; Kumar et al., 2010), encode memories (Robinson et al., 2017), or manage decision conflicts (Smith et al., 2019). The most well-known neural coding scheme is the single unit firing rate (Gerstner et al., 1997), or ‘rate coding,’ which is studied extensively to determine how firing rates support important functions such as perception (Andrei et al., 2019) and motor control (Kline and De Luca, 2015). Rate coding and temporal coding are potentially independent mechanisms that can occur within the same region (Andrei et al., 2019; Gerstner et al., 1997), but both are affected by the oscillating FP (Buzsáki et al., 2012; Rasch et al., 2008). The SFC results suggest that distant field potentials occurring several millimeters away can be weakly coupled to local spike timing in the human neocortex. Our findings are consistent with previous work showing that local unit activity can be coherent with low frequency oscillations measured at a distance (Benchenane et al., 2010; Eissa et al., 2017).

This research expands our knowledge on the effective range of phase coherence and spike-field coupling across the cortical mantle. The spatial reach of coherence seems similar across brain regions, but it is likely that more complex spatial patterns of coherence exist when examined across different cortical and subcortical structures. We encourage the use of simultaneous ECoG, depth electrode, and/or MEA recordings to further advance our knowledge on the spread of information through cerebral space. With these ideas in mind, future studies can include more subjects, more brain regions, and increase focus on brain state or task-related changes.

## METHODS

### Subjects

Three neurosurgical patients (3 male, mean age = 28 y/o) with medically refractory epilepsy underwent craniotomies for the placement of subdural ECoG grids in frontocentral cortex (subject A), middle temporal cortex (subject B), and posterior temporal cortex (subject C). Decisions regarding the location and coverage of the ECoG arrays were based solely on clinical criteria. A 96-channel Utah microelectrode array (UMA) was placed underneath the ECoG grid for the purpose of recording seizures. The Columbia University Medical Center Institutional Review Board approved placement of these FDA-approved research electrodes in regions that are likely to be part of the eventual resection (IRB-AAAB6324). All patients provided informed consent before participating in the study.

### Electrophysiological Recordings and Preprocessing

The ECoG and the UMA data were recorded from the frontal cortex, the middle temporal cortex and posterior temporal cortex. A neural signal processor (Blackrock Microsystems) simultaneously acquired the ECoG and UMA data, which were amplified, band-pass filtered (0.3 Hz – 7.5 kHz), and digitized at 2kHz and 30kHz, respectively. All ECoG data were re-referenced into a bipolar montage. Similarly, bipolar derivations from the UMA were used as a representation of deep cortical field potentials. Behavioral trials that contained epileptiform discharges were removed based on visual inspection and/or aberrant voltage deflections (*μ*V) greater than 5 standard deviations across trials.

### Time-frequency Decomposition and Neuronal Coherence

Field potential spectra were computed via Fast Fourier Transform (FFT) with a Hanning window taper (window size: 256 ms) using EEGLAB functions and custom code written in MATLAB (Delorme and Makeig, 2004). The frequency bands of interest were delta (1-3 Hz), theta (4-7 Hz), alpha (8-14 Hz), beta (15-30 Hz), and gamma signals (31-50 Hz). Instantaneous FP phase was extracted by taking the angle between the real and imaginary components of the spectral decompositions. As a measure of neuronal coherence, we used the phase lag index (PLI), which represents the consistency of nonzero phase angle differences between two time series (Stam et al., 2007). The PLI is computed as *PLI*(*t,f*)=|⟨*sign*|Δ _Φ_(*t*_*k*_)|⟩|, where Δ _Φ_(*t*_*k*_), where Δ _Φ_(*t*_*k*_)is the instantaneous phase difference between two time series at time *t, k* = 1 … N, for each frequency, *f*. The signum operation within the PLI reduces its sensitivity to volume conduction because instantaneous coupling is removed. A PLI of 1.0 implies perfect phase alignment, whereas a PLI of 0.0 implies an absence of coupling. ‘Surface coherence’ was computed as the PLI between the ECoG channel closest to the UMA, and all other ECoG channels (0.5 - 1 cm spacing). ‘Surface-depth coherence’ was computed as the PLI between the superficial ECoG and deep cortical FP (MEA). The PLI is based entirely on the phase component of neural signals, making it resistant to spectral power fluctuations that can affect signal-to-noise ratios. We were exclusively interested in phase synchrony between neural signals, as the relationship between power and phase during cognitive tasks has been dissociated in some studies (van Diepen et al., 2015). The average difference in spectral power (1-50 Hz) between ECoG and deep cortical FPs was only 1.14 ± 0.21 dB (Figure S3). For each subject, PLI was averaged across all behavioral trials for each electrode.

### Action Potential Sorting

Single unit activity (SUA) data was thresholded at -4 times the root mean square of the high-pass filtered signal (> 250 Hz). The action potential waveforms were then separated based on t-distribution expectation maximization (EM) computed on a feature space comprised of the first three principal components (Shoham et al., 2003), using Offline Sorter (Plexon, inc). Spike sorting yielded well-isolated neurons from the frontocentral cortex (n = 100), middle temporal cortex (n = 191), and posterior temporal cortex (n = 206).

### Spike-field Coherence

Spike-field coherence (SFC) was computed via multi-taper approach using the Chronux toolbox for MATLAB (Mitra and Pesaran, 1999). The SFC computation included a 10 ms step size, a time-bandwidth product of five, and a sequence of nine discrete prolate spheroidal (dpss) tapers. Multi-taper approaches have been shown to improve consecutive spectral estimates, which is useful for SFC analyses (Hoogenboom et al., 2006). The multi-taper estimate of a spike train’s spectrum can be calculated as,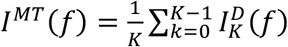, where the eigenspectra 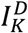are direct spectral estimates obtained from *k*_*n*_ taper functions for the duration of samples, *D* (Jarvis & Mitra, 2001). SFC was computed as the magnitude squared coherence between the multi-taper spectral estimate of the spike train and the concurrent FP (Baker et al., 2003; Zeitler, Fries, & Gielen, 2006). For each subject, SFC was averaged across all neurons and behavioral conditions for each electrode.

### Spatial Distance Modeling

Coherence (PLI or SFC) was modeled as a function of Euclidean distance (mm) between the ECoG contact closest to the MEA and all other ECoG contacts on the grid. The MEA location was chosen as the origin point for the distance calculations because it allowed for comparing the lateral spread of ECoG-, deep cortical-, and spike-field coherence across the same geometric space. The 3-dimensional coordinates for all contacts were extracted via co-registration of preoperative structural MRI with postoperative CT images using the Statistical Parametric Mapping (SPM) Toolbox. A single power function was fit to the mean coherence values for each channel’s distance from the origin. The general form of the function is given by, *Coherence*=*a*\**disItance*^*b*^, where *a* and *b* are model coefficients. Nonlinear regression was performed using the ‘curve fitting toolbox’ in MATLAB. We used a power function because it is theoretically aligned with previous work on signal propagation in neural systems (Klaus et al., 2011). The function allows for the estimation of coherence if one knows the distance from a spatial origin point. The goodness-of-fit metric for the functions is given by their *r*^*2*^ value, which represents the magnitude of the correlations between the raw and model-predicted values of coherence as a function of distance. Comparisons with other functions such as linear, exponential, and Fourier series were not of interest in this study. The spatial reach of coherence was estimated via changepoint detection (CPD) using the ‘findchangepts’ function in MATLAB (Killick et al., 2012, Lavielle, 2006).

Changepoints were detected using a log likelihood approach based on the variance of the standard deviation of coherence values, *x*, given by 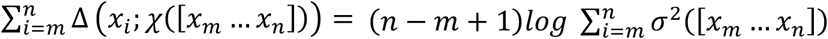, where χ is the empirical estimate and Δ is the deviation measure. Therefore the spatial reach of coherence was defined as the average physical distance in millimeters where the variance of coherence’s standard deviation changed significantly. Normalization via median centering was applied to the coherence data so that spatial reach across all brain regions could be estimated using values on a scale relative to their own median coherence.

### Cognitive Control Task

The first subject (A) had frontocentral ECoG coverage and performed 3 sessions of the multi-source interference task (MSIT) (Bush and Shin, 2006). MSIT combines two different types of conflict associated with cognitive control, known as ‘spatial’ & ‘distractor’ conflict. Each trial began with a 500 ms visual fixation period, followed by a cue that consisted of three numbers that ranged from 0 to 3 (white color, black background). The ‘target’ number was one of the three numbers that was different than the other two ‘distractor’ numbers. Patient 1 used a 3-button response pad to indicate which number was the target (number 1: left-hand button, number 2: middle button, number 3: right-hand button). Spatial conflict occurred if the target number was in a different position than its representation on the response pad (e.g., ‘3 0 0’; the target is in the left-hand position, but the right-hand button is the correct choice). Distractor conflict occurred if the distractor numbers were possible button choices (e.g., ‘1 3 1’, where ‘1’ corresponds to a possible button choice). Some trials contained neither type of conflict (e.g., ‘0 2 0’), whereas others contained both conflict types (e.g., ‘2 1 1’). After the patient responded on each trial, the cue disappeared, and feedback appeared with variable delay (300-800 ms). In blocks of ten trials, the feedback consisted of the target number in a different color (green for correct, red for incorrect), alternating with neutral feedback that was blue, regardless of accuracy. The inter-trial interval varied uniformly randomly between 1000 ms and 1500 ms. Across three sessions, Patient 1 performed 713 trials. To maintain simultaneity of the single unit recordings across patients, SFC data was used only for the first session, which was comprised of 295 trials. All neural data for this task were epoched from 500 milliseconds (ms) before until 1500 ms after the time of cue onset.

### Speech Perception Task

Subject B had middle temporal ECoG coverage and they listened to a series of audio clips of male and female voice actors, where both voice actors read different stories at the same time. The audio clips were played from a single device in front of the subject (Bose Soundlink Mini). The task was split into 4 blocks, where the subject was asked to attend to either the male or female speaker, alternating between each block. During intermittent pauses, the subject was asked to repeat the last sentence of the attended speaker. The neural data were epoched from 1000 ms before until 5000 ms after the time of stimulus onset.

### Social Cognition Task

Subject C had posterior temporal ECoG coverage and watched several repetitions of 18 unique video clips from popular films (e.g. ‘Girls Trip’, ‘Split’, ‘Get Out’, and ‘The Hours’) in which a salient emotion was displayed, and a full sentence was spoken (e.g. “the chores have become my sanctuary.”). Each trial began with a visual fixation period (500 ms), followed by the video clip. The neural data were epoched from 1000 ms before until 5000 ms after the time of video onset.

## QUANTIFICATION AND STATISTICAL ANALYSIS

All curve fitting procedures were performed in MATLAB (MathWorks). All group-level statistical analyses were performed in SPSS (IBM). Mixed ANOVAs were used throughout, which utilized a 2 (*distance*: proximal, distal) x 3 (depth: surface, surface-depth, SFC) x 5 (*frequency*: delta, theta, alpha, beta, gamma) design. Bonferroni correction was used to correct for multiple comparisons and the Greenhouse-Geiser method was used to adjust degrees of freedom when sphericity assumptions were violated. Box and whiskers are inter-quartile range (IQR) and 1.5 x IQR respectively, and asterisks represent outliers. Correlations between neuronal coherence and spectral power were computed using Pearson’s *r* (see Figures S4-S6). Statistical tests used in this study are also described in the figure legends.

## Acknowledgements

This study was supported by R01 MH106700 (SAS), Dana Foundation (SAS), McNair Foundation (SAS) and the Brain & Behavior Research Foundation (#26706; EHS).

## AUTHOR CONTRIBUTIONS

Conceptualization: JM, ES, SS

Methodology: JM, ES

Investigation: ES, JO

Software: JM, ES

Formal Analysis: JM, ES

Writing – Original Draft: JM

Writing – Review & Editing: JM, ES, ML, CS, SS

Funding Acquisition: SS

---

**Figure S1.**
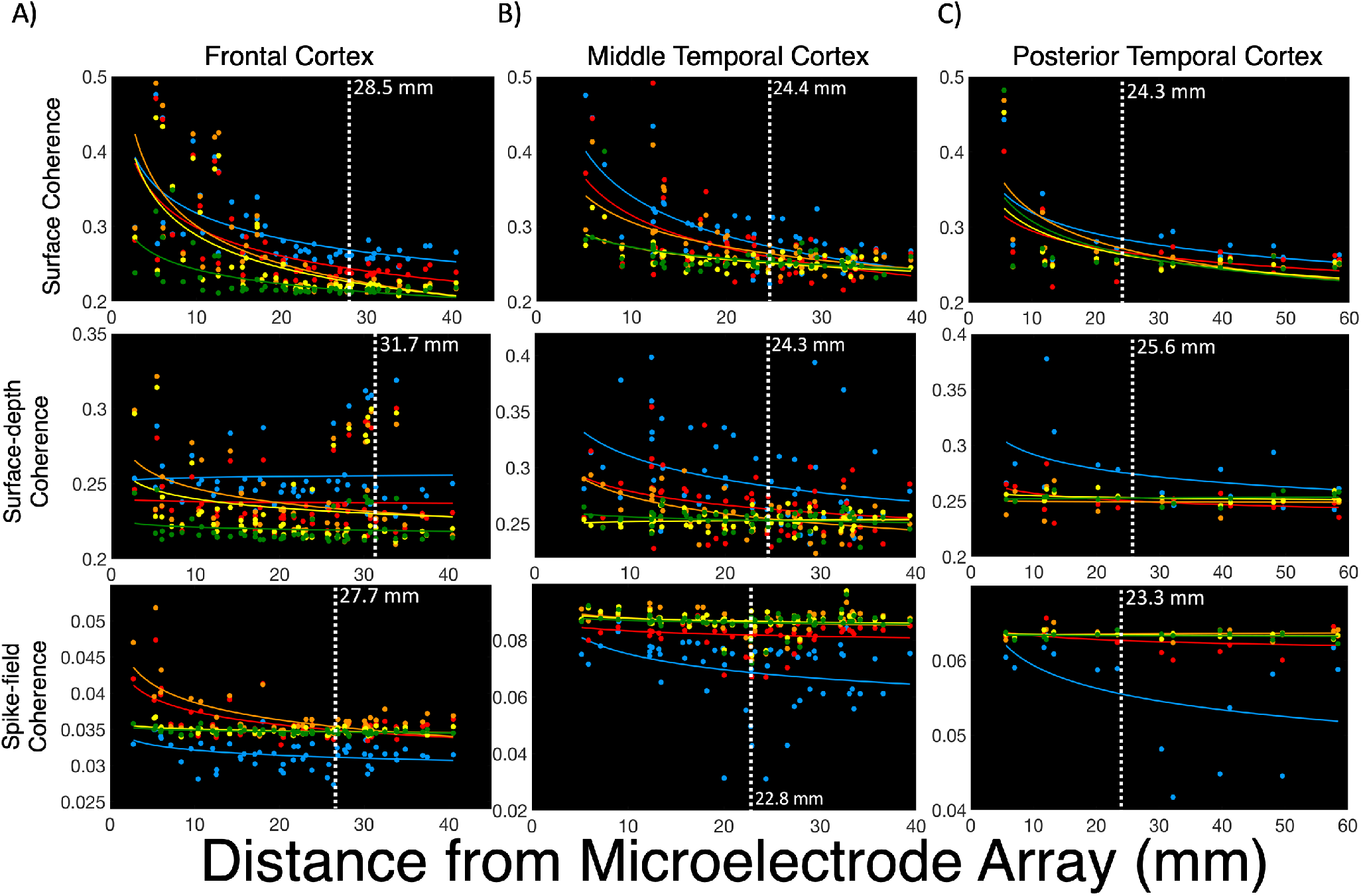
Spatial Reach is Similar across Brain Regions. Across frontocentral cortex (A), surface coherence reached 2.84 cm, showing significant power law decay for delta (*p* < 0.001), theta (*p* < 0.001), alpha (*p* < 0.001), beta (*p* < 0.001), and gamma (*p* < 0.001) signals. Surface-depth coherence reached 3.17 cm across frontocentral cortex showed decay for only alpha (*p* = 0.027) signals. SFC reached an estimated 2.77 cm across the frontocentral cortex. Power law decay was observed for delta (*p* = 0.011), theta (*p* < 0.001), alpha (*p* < 0.001), beta (*p* = 0.003), and gamma (*p* = 0.001) SFC. Across middle temporal cortex (B), surface coherence reached approximately 2.44 cm, showing decay for delta (*p* < 0.001), theta (*p* < 0.001), alpha (*p* < 0.001), beta (*p* < 0.001), and gamma (*p* < 0.001) signals. Surface-depth coherence reached 2.43 cm across middle temporal cortex, showing decay for delta (*p* = 0.008), theta (*p* = 0.022), and alpha (*p* < 0.001) signals. Spike-field coherence (SFC) across the middle temporal cortex reached an estimated 2.27 cm across temporal cortex, showing decay was observed for delta (*p* = 0.005), theta (*p* = 0.004), alpha (*p* = 0.001), beta (*p* = 0.001) and gamma SFC (*p* = 0.001). Across posterior temporal cortex (C), surface coherence reached 2.43 cm and showed decay for delta (*p* = 0.020), theta (*p* = 0.043), alpha (*p* = 0.043), beta (*p* = 0.031), and gamma (*p* = 0.025) frequencies. Surface-depth coherence reached 2.56 cm, and showed no showed no power law decay across posterior temporal cortex. Spike-field coherence (SFC) reached an estimated 2.21 cm. There was no significant decay for SFC in this region, and only delta signals showed a trend (*p* = 0.087).

**Figure S2.**
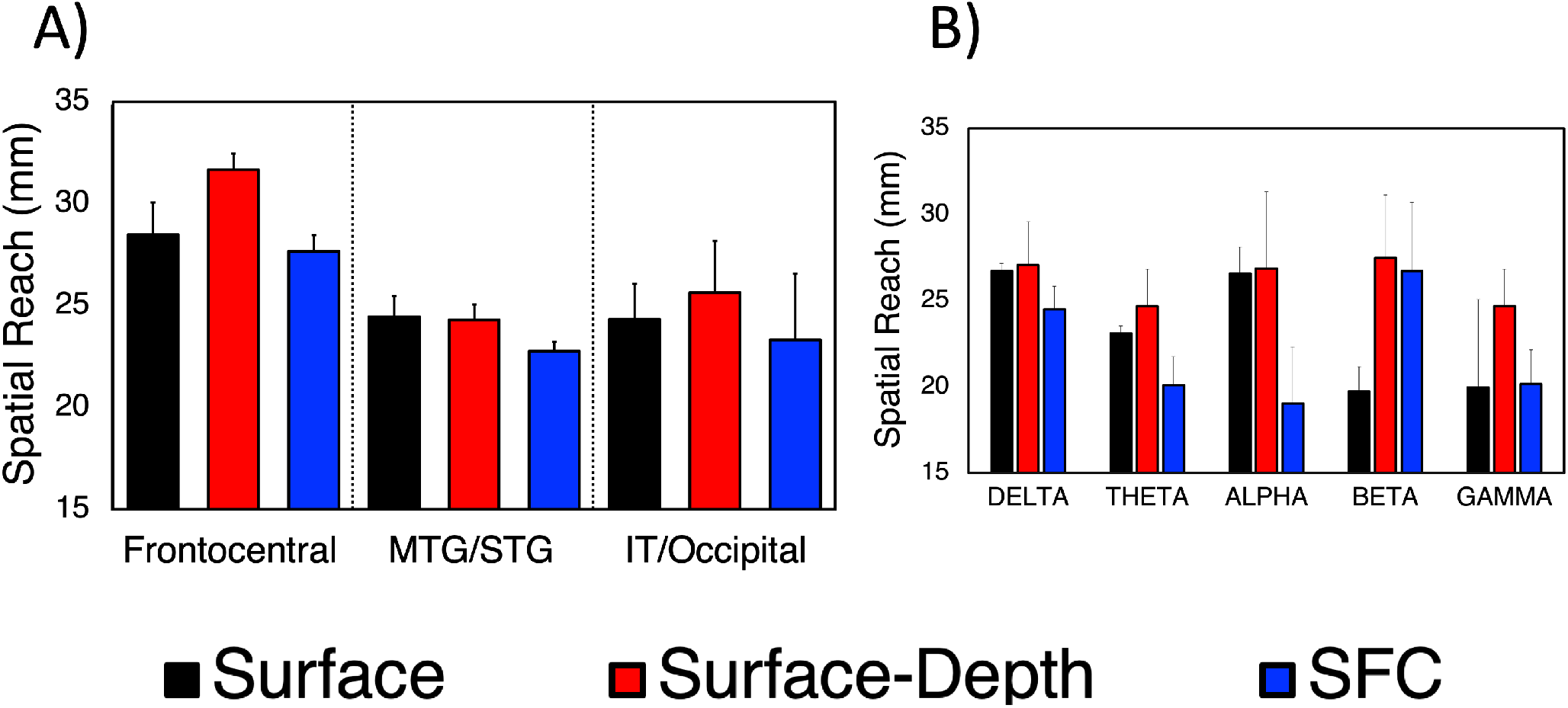
Spatial Reach of Coherence is Similar across Brain Regions and Frequencies. The spatial reach of neuronal coherence and SFC was significantly greater for frontal cortex compared to other brain regions (*η*^*2*^ = 0.82, p *<* 0.001) (A). Although surface-depth coherence appeared to reach farther than surface coherence and SFC, we did not observe a significant effect of depth (p *=* 0.178) or frequency (p *=* 0.869) (B). Across all brain regions and types of coherence, the spatial reach of coherence was approximately 2.58 cm (±1.45 mm). The spatial reach of coherence was relatively stable across brain regions and frequencies. These results were computed using ANOVAs (depth(3) x region(3); depth(3) x frequency(5)).

**Figure S3.**
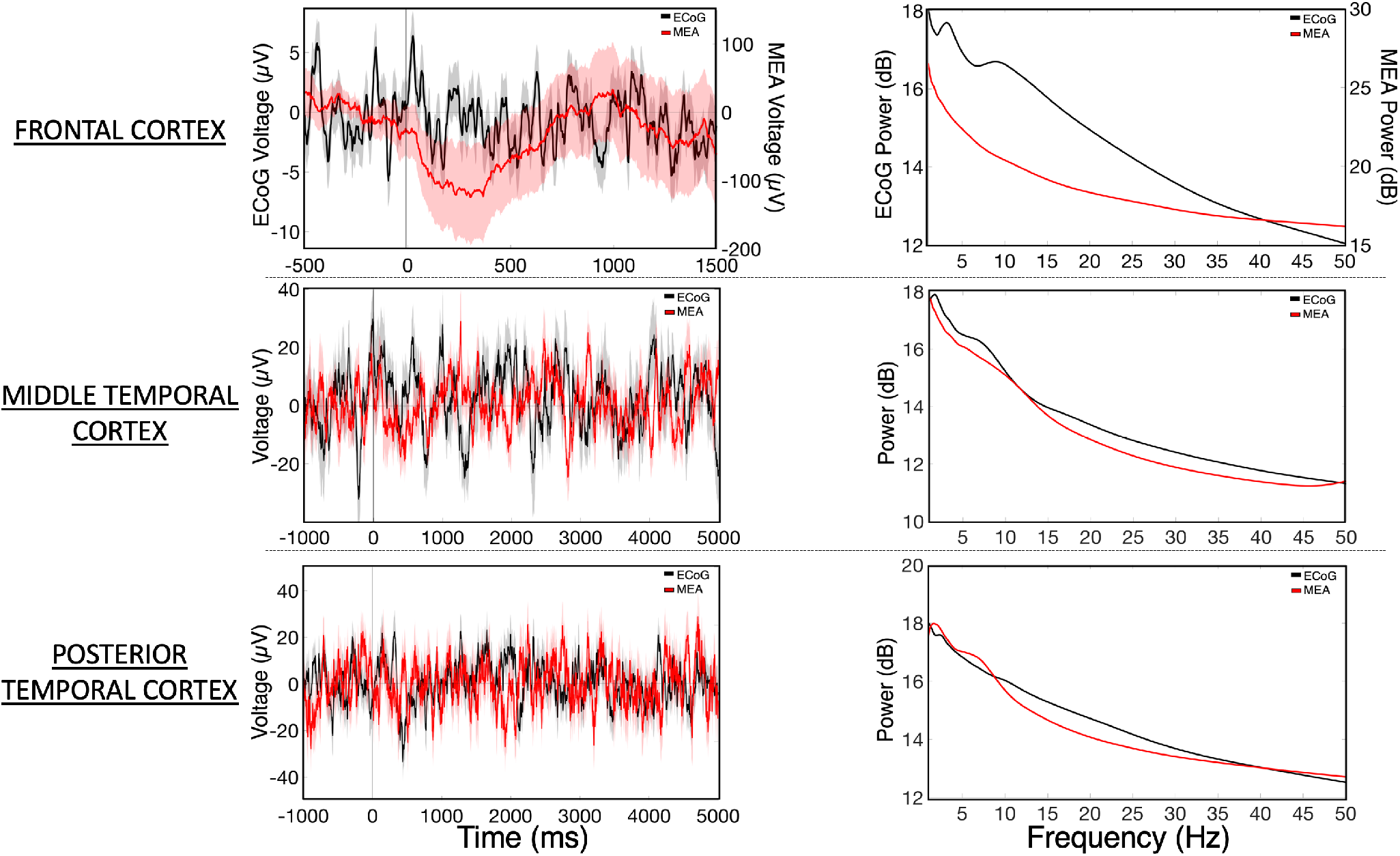
ECoG vs LFP Voltage Traces and Power Spectra. The average difference in spectral power (1-50 Hz) between ECoG and deep cortical field potentials recorded from microelectrode arrays was 1.14 ± 0.21 dB. The variance in power for deep cortical field potentials encouraged the use of PLI to evaluate phase synchrony. Since PLIs are based solely on the phase angle differences between separate neural signals, the coherence results are less sensitive to changes in spectral power. Shaded regions depict standard error of the mean across subjects.

**Figure S4.**
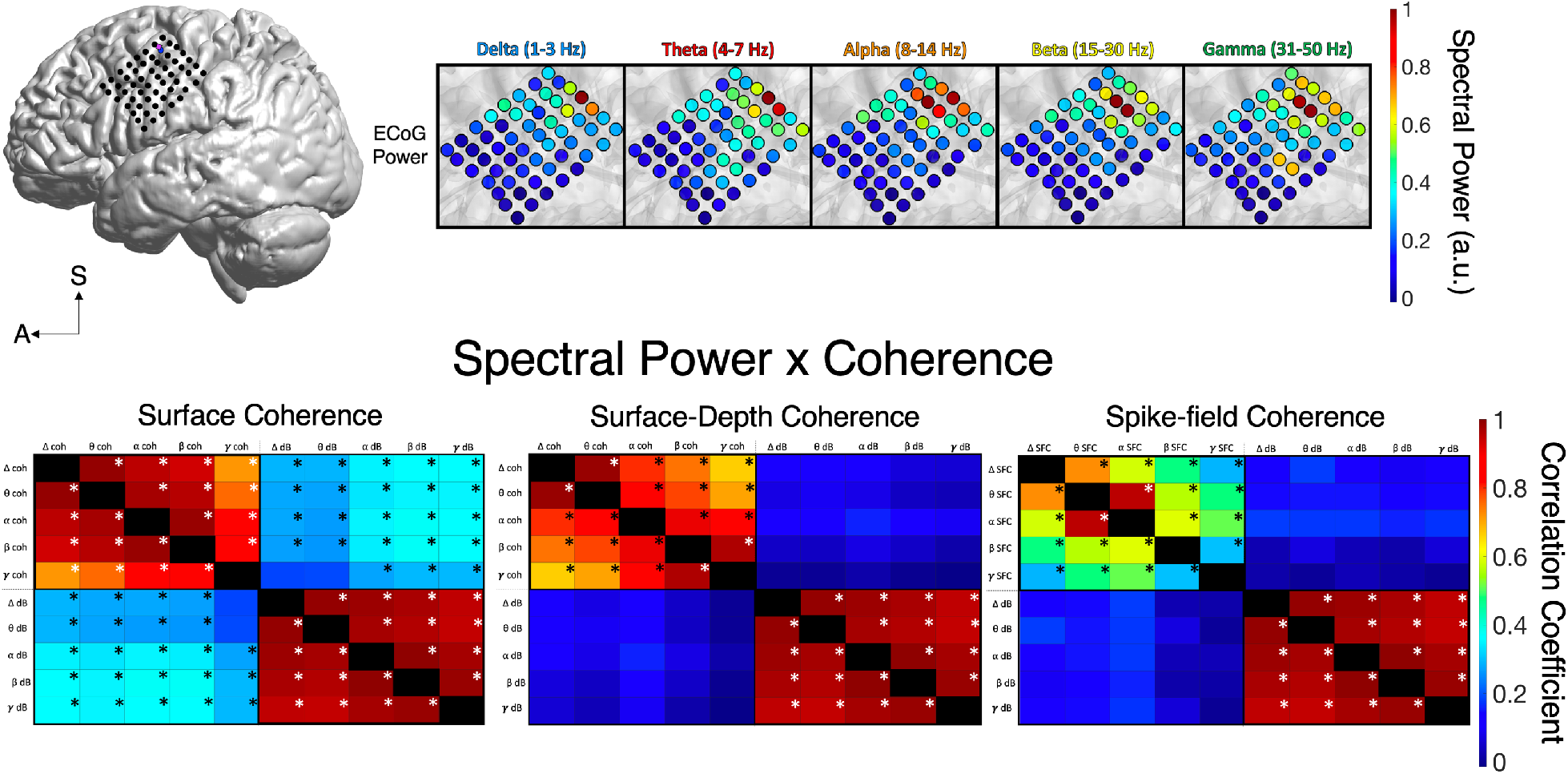
Correlations between Coherence and Spectral Power across Frontal Cortex. Correlations between spectral power and coherence. Spectral power was moderately correlated with surface coherence across frontal cortex. Surface-depth and spike-field coherence did no show significant correlations with spectral power. Greek symbols refer to delta (1-3 Hz), theta (4-7 Hz), alpha (8-14 Hz), beta (15-30 Hz), and gamma (31-50 Hz) frequency bands. Asterisks indicate statistically significant correlations *(p* < 0.05).

**Figure S5.**
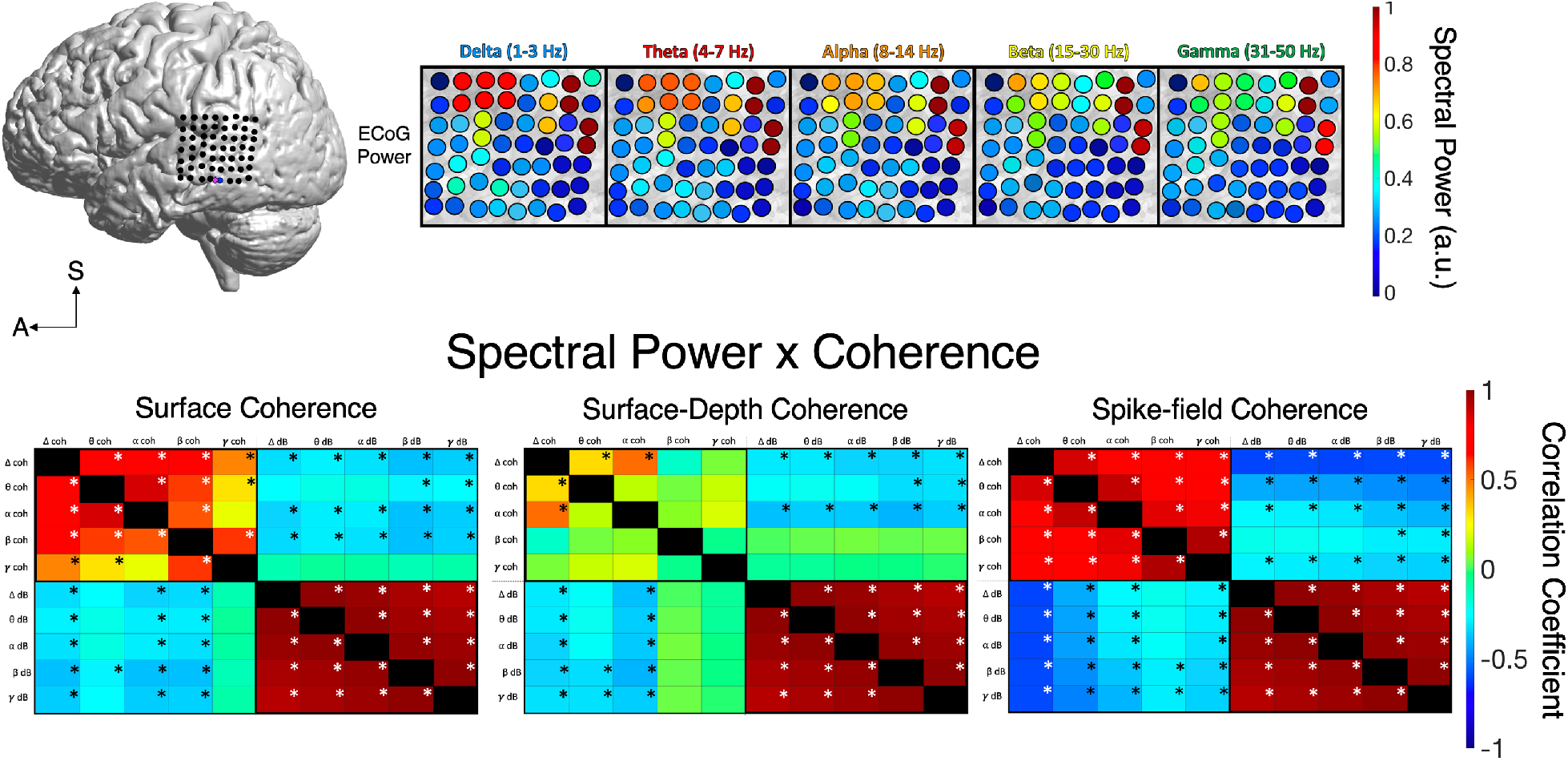
Correlations between Coherence and Spectral Power across Middle Temporal Cortex. Correlations between spectral power and coherence. Spectral power was negatively correlated with coherence across middle temporal cortex. Greek symbols refer to delta (1-3 Hz), theta (4-7 Hz), alpha (8-14 Hz), beta (15-30 Hz), and gamma (31-50 Hz) frequency bands. Asterisks indicate statistically significant correlations (*p* < 0.05).

**Figure S6.**
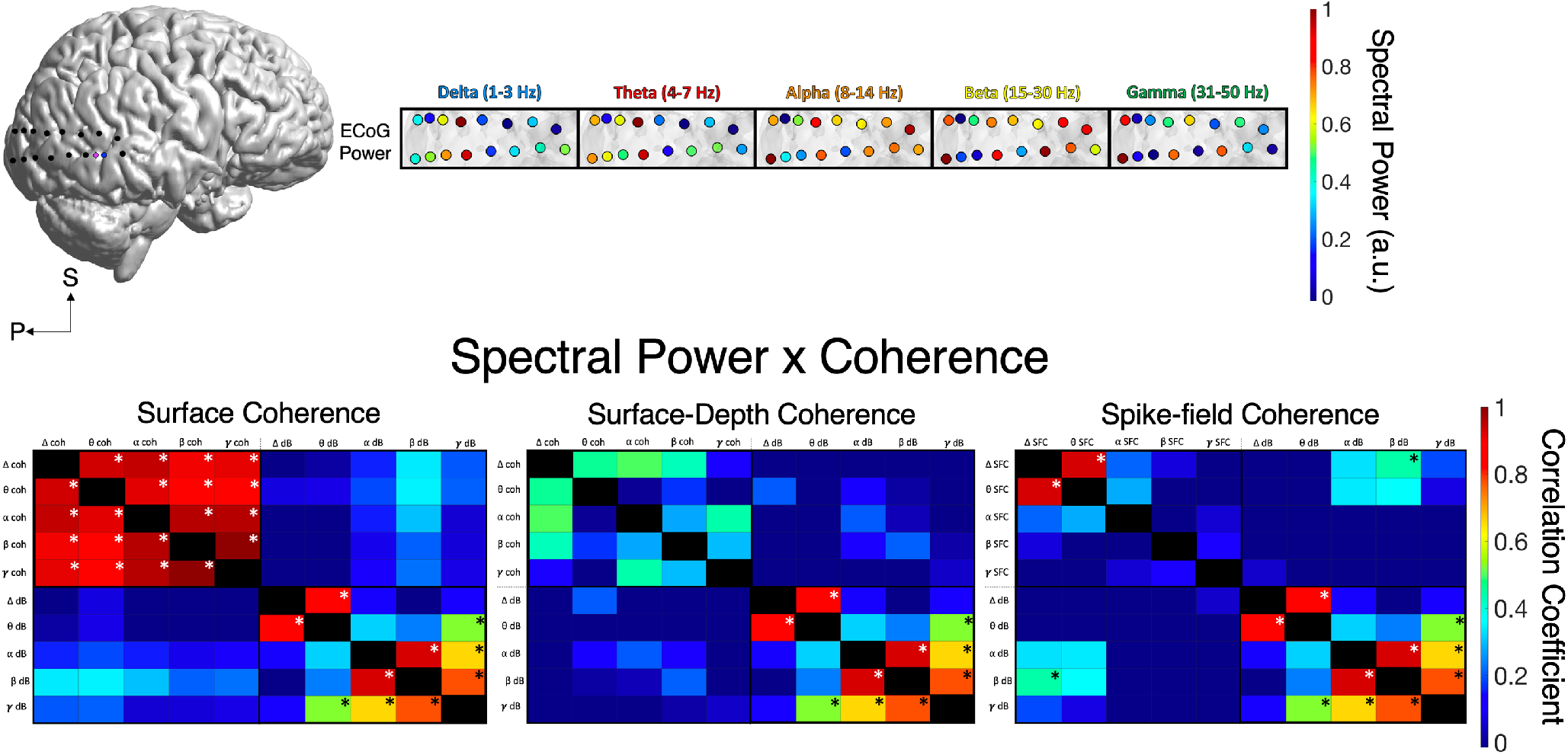
Correlations between Coherence and Spectral Power across Posterior Temporal Cortex. Correlations between spectral power and coherence. Spectral power was moderately correlated with beta (15-30 Hz) spike-field coherence across posterior temporal cortex. Surface- and surface-depth coherence did no show significant correlations with spectral power. Greek symbols refer to delta (1-3 Hz), theta (4-7 Hz), alpha (8-14 Hz), beta (15-30 Hz), and gamma (31-50 Hz) frequency bands. Asterisks indicate statistically significant correlations (*p* < 0.05).

## Notes

### Competing Interest Statement

The authors have declared no competing interest.

